# The effect of cytoskeleton inhibitors on coccolith morphology in *Coccolithus braarudii* and *Scyphosphaera apsteinii*

**DOI:** 10.1101/2022.02.18.480830

**Authors:** Gerald Langer, Ian Probert, Alison Taylor, Glenn M. Harper, Colin Brownlee, Glen Wheeler

## Abstract

The calcite platelets of coccolithophores (Haptophyta), the coccoliths, are among the most elaborate biomineral structures. How these unicellular algae accomplish the complex morphogenesis of coccoliths is still largely unknown. It has long been proposed that the cytoskeleton plays a central role in shaping the growing coccoliths. Previous studies have indicated that disruption of the microtubule network led to defects in coccolith morphogenesis in *Emiliania huxleyi* and *Coccolithus braarudii*. Disruption of the actin network also led to defects in coccolith morphology in *E. huxleyi*, but its impact on coccolith morphology in *C. braarudii* was less clear, as coccolith secretion was largely inhibited under the conditions used. A more detailed examination of the role of actin and microtubule networks is therefore required to address the wider role of the cytoskeleton in coccolith morphogenesis. In this study, we have examined coccolith morphology in *C. braarudii* and *Scyphosphaera apsteinii* following treatment with the microtubule inhibitors vinblastine and colchicine (*S. apsteinii* only) and the actin inhibitor cytochalasin B. We found that all cytoskeleton inhibitors induced coccolith malformations, strongly suggesting that both microtubules and actin filaments are instrumental in morphogenesis. By demonstrating the requirement for the microtubule and actin networks in coccolith morphogenesis in diverse species, our results suggest that both of these cytoskeletal elements are likely to play conserved roles in defining coccolith morphology.

## Introduction

Coccolithophores, haptophyte algae, are among the most important pelagic calcite producers (Baumann et al. 2004, Poulton et al., 2007, Ziveri et al., 2007). The calcite platelets (coccoliths) that form the cell covering display an intricate morphology including elaborately shaped crystals in the diploid life cycle stage (Young et al. 1999). Although definitive evidence for the precise function of calcification in coccolithophores has been difficult to obtain, it is likely that assembly of coccoliths into a protective coccosphere is central to their function (Monteiro et al 2016). For instance, it was shown that the interlocking coccosphere of *E. huxleyi* confers remarkable mechanical protection, and *C. braarudii* needs an intact coccosphere to divide (Jaya et al 2016, Walker et al 2018). The distinct, normal morphology of the coccoliths is required for the correct formation of the coccosphere (Young 1994, Henriksen et al. 2003, Bown et al 2004, Quintero-Torres et al 2006, Jaya et al. 2016, Walker et al. 2018). Morphogenesis of coccoliths is therefore a central element of coccolithophore eco-physiology and evolution. Despite this prime position in coccolithophore biology, the morphogenesis of coccoliths is not well understood. Just over a decade ago coccolith morphogenesis was still regarded as “the most enigmatic part of biomineralization” (Henriksen et al 2004, p. 726). Although some progress has been made in the last decade (see below), this statement has lost little of its edge.

While Huxley (1868) regarded coccoliths as of inorganic origin, it is now clear that the morphologies of coccolith crystals are not to be found in inorganically precipitated calcite (Young et al. 1999, Aquilano et al. 2016). Calcification in coccolithophores occurs intracellularly, allowing precise control of the chemical conditions for the precipitation of calcium carbonate. The coccolith develops in a specialised intracellular compartment, the coccolith vesicle (Dixon 1900, Wilbur and Watabe 1963), where calcium carbonate crystals are nucleated onto an organic baseplate to produce small, initially rhombic, crystals. The calcite crystals then undergo carefully controlled growth to produce mature coccoliths with distinctive morphologies for each species. The mature coccoliths are subsequently secreted to the cell surface, where they are arranged to form the coccosphere. The cytoskeleton likely plays several important roles in coccolithogenesis, including controlling the secretion of the coccolith vesicle to the cell surface. Significant research interest has focused on the requirement for the cytoskeleton in shaping the morphology of the developing coccolith.

The coccolith vesicle adopts the shape of the growing coccolith (Outka and Williams 1971, Klaveness 1972, Westbroek et al. 1984, Probert et al. 2007), which has led to the hypothesis that the coccolith vesicle acts as dynamic mould for the developing coccolith (Klaveness 1972, 1976, Young et al. 1999). This view includes a controlled force that shapes the coccolith vesicle. Based on transmission electron microscopy (TEM) examination of developing coccoliths it was hypothesised that a fibrillar structure adjacent to the coccolith vesicle exerts this force (Klaveness 1972, 1976). This led to the idea that the fibrillar material, later equated with the cytoskeleton (Remak 1843, Freud 1882), is at the centre of the coccolith shaping machinery (Westbroek et al. 1984, Didymus et al. 1994, Marsh 1994, 1999, Young et al. 1999, 2009, Marsh et al. 2002). Although this hypothesis is widely accepted, the supporting evidence from TEM analysis remains somewhat ambiguous (Klaveness et al 1972, 1976). This ambiguity was not eliminated by later TEM studies (Westbroek et al. 1984, Taylor et al. 2007). Recently immunofluorescence microscopy has revealed a microtubule network in close contact with the coccolith vesicle in *C. braarudii* (Durak et al. 2017). This observation complements the earlier TEM studies and strongly supports the original dynamic mould hypothesis (Klaveness 1972, 1976).

The cytoskeleton is central to many aspects of cellular function. Whilst many chemical inhibitors exist that disrupt the function of the microtubule and actin networks within the cell, their use to examine specific processes is complicated by their potential to interfere with other aspects of cell physiology. However, Langer et al 2010 demonstrated that the application of microtubule and actin inhibitors to coccolithophores could be carefully titrated to partially disrupt cytoskeleton function without complete inhibition of cellular growth. Application of the microtubule inhibitor colchicine or the actin inhibitor cytochalasin B to the abundant bloom-forming species *Emiliania huxleyi* resulted in significant disruption of coccolith morphology. These malformations were not observed in other treatments that reduced growth rate (e.g. the photosynthesis inhibitor 3-(3,4-dichlorophenyl)-1,1-dimethylurea (DCMU)), leading to the conclusion that both actin and microtubules play a central role in controlling the morphology of the developing coccolith. These findings therefore provide experimental support for the dynamic mould hypothesis.

A subsequent study found similar effects on coccolith morphology in *Coccolithus braarudii* using the microtubule inhibitor nocodazole (Durak et al. 2017). The effect of disrupting actin in *C. braarudii* was however different, as it led to a complete inhibition of coccolith production. It is therefore possible that actin is not involved directly in coccolith morphogenesis in *C. braarudii* but plays a more general role in coccolith production, such as the exocytosis of coccoliths, or that actin is needed to start the whole process of coccolith formation (see Durak et al. 2017). Considering that both TEM and immunofluorescence imaging (Taylor et al. 2007, Durak et al. 2017) have so far only provided evidence for the involvement of microtubules in morphogenesis, evidence for the role for actin in coccolithogenesis remains limited.

It is important to note that the pharmacological agents used to disrupt the cytoskeleton in these studies have distinct modes of action (Supplementary Table 1) (Langer 2010; Durak 2017). Moreover, Langer et al 2010 examined coccolith morphology in *E. huxleyi* cells grown in test conditions for several generations, whereas Durak et al 2017 disrupted cytoskeletal networks in decalcified *C. braarudii* cells and then assessed their ability to recalcify. These methodological differences make it difficult to directly ascertain the wider requirement for actin in coccolith formation in coccolithophores.

We have therefore performed a detailed examination of the impact of cytoskeleton disruption on coccolith formation in two coccolithophore species. *C. braarudii* is a heavily calcified species that is abundant in temperate and sub-polar regions of the North Atlantic and contributes significantly to calcification in this regions (Daniels et al 2016). To obtain a broader picture of the effects of cytoskeleton inhibitors on coccolith morphology, we have additionally examined *Scyphosphaera apsteinii*. This dimorphic species produces two distinct coccolith types, the disc-like muroliths and the large barrel-shaped lopadoliths (Drescher 2012). Moreover, *S. apsteinii* is a member of the Zygodiscales and therefore occupies a distinct evolutionary lineage from *E. huxleyi* (Isochrysidales) and *C. braarudii* (Coccolithales). We have treated *C. braarudii* and *S. apsteinii* with a range of inhibitors that act to disrupt actin and microtubule function within the cell. We show that both components of the cytoskeleton play an important role in coccolith morphogenesis in these species.

## Material and Methods

### Culture conditions

Clonal cultures of *C. braarudii* (strain PLY182g) and *S. apsteinii* (strain RCC1456) were grown in aged (3 months), sterile-filtered (Stericup-GP Sterile Vacuum Filtration System, 0.22 μm pore size, polyethersulfone membrane, Merck) natural surface seawater sampled in the English Channel off Plymouth, UK (station E1: 50°02.00′ N, 4° 22.00′ W) enriched with 100 μM nitrate, 6.25 μM phosphate, 4 μM silicate, 0.005 μM H2SeO3, 0.00314 μM NiCl2, and trace metals and vitamins as in f/2 medium (Guillard 1975). Strain RCC1456 was obtained from the Roscoff Culture Collection (http://www.sb-roscoff.fr/Phyto/RCC), and strain PLY182g from the Plymouth Culture Collection (https://www.mba.ac.uk/facilities/culture-collection#b7).

Cultures were grown under a 16:8 light:dark cycle. Experiments were carried out at a light intensity of 50 μmol photons m^-2^ s^-1^ in temperature controlled culture rooms. *C. braarudii* PLY182g was grown at 15 °C, and *S. apsteinii* was grown at 18 °C. Cells were grown in dilute batch cultures, ensuring a quasi-constant seawater carbonate system over the course of the experiment (Langer et al. 2013). Each data point is the mean value of triplicate culture experiments. Error bars represent SD.

For determination of cell density, samples were taken every other day (or less frequently, depending on growth rate) and counted immediately after sampling using a Sedgwick Rafter Counting Cell. Cell densities were plotted versus time, and growth rate (μ) was calculated from exponential regression using the natural logarithm.

### Application of cytoskeleton inhibitors

Colchicine, vinblastine, and cytochalasin B were obtained from Sigma-Aldrich (Munich, Germany).

Vinblastine was dissolved in reverse osmosis water. The concentration of the stock solution was 1.1 mM. *C. braarudii* was treated with a final vinblastine concentration of 2 μM, and *S. apsteinii* with a final vinblastine concentration of 1.25 μM.

Colchicine was dissolved in culture medium. The concentration of the stock solution was 2.5 mM. *S. apsteinii* was treated with a final colchicine concentration of 20 μM.

Cytochalasin B stock solution (20.9 mM in DMSO) was obtained from Sigma-Aldrich (Munich, Germany).*C. braarudii* was treated with a final cytochalasin B concentration of 1.5 μM, and *S. apsteinii* with a final cytochalasin B concentration of 1 μM. Consequently, cells were exposed to a maximum DMSO concentration of 0.007 vol %. This DMSO concentration is harmless; it was shown that in *E. huxleyi* 0.5 vol % DMSO has no effect on growth rate (Langer et al. 2010). In confirmation *C. braarudii* and *S. apsteinii* grown in 0.01 vol % DMSO showed normal growth and, upon qualitative inspection by means of light microscopy, no notable increase in coccolith malformations.

All stock solutions were freshly prepared prior to the start of the experiments. Cells were exposed to cytoskeleton inhibitors for 25 d, after which samples were taken for analysis of coccolith morphology.

### SEM analysis of coccolith morphology

Samples for SEM analysis were filtered on polycarbonate filters (0.8 μm pore-size), dried in a drying cabinet at 50°C for 24 h, then sputter-coated with gold-palladium using an Emitech K550 sputter coater at Plymouth Electron Microscopy Centre (PEMC). Imaging was performed with both Jeol JSM-6610LV and Jeol JSM-7001F at PEMC. The following categories were used to describe coccolith morphology. 1) *C. braarudii*: normal, malformation type R, minor malformation, major malformation, rhomb-like malformation. For reference images see Fig. 1. A preliminary analysis showed that the percentage of incomplete coccoliths was less than 1 % in all samples. Therefore, incomplete coccoliths were not accurately quantified in the final analysis. 2) *S. apsteinii:* normal, malformation type R, malformation type S, malformation type T. For reference images see Fig. 5. An average of ~350 coccoliths was analysed per sample (Langer and Benner 2009). The methodology of coccolith categorization and counting employed here is well established and yields robust and unbiased results (Langer et al. 2006; Langer and Benner 2009; Langer and Bode 2011; Langer et al. 2011; Langer et al. 2012, Bach et al. 2012). The percentage of intact, as opposed to collapsed, coccospheres was analysed in the same samples as coccolith morphology. An average of ~300 coccospheres was analysed per sample. Data presented in the figures are averages of triplicate cultures; error bars represent SD. The percentage of intact coccospheres was only analysed in *C. braarudii* because *S. apsteinii* coccoliths do not interlock and show high percentages of collapsed coccospheres in all samples. Please note that coccospheres that collapse during preparation for SEM imaging could well have been perfectly intact in culture. Preparation for SEM imaging imposes mechanical stress on coccospheres that often leads to the collapse of non-interlocking coccospheres such as those of *S. apsteinii*. By contrast, interlocking coccospheres have exceptional mechanical stability (Jaya et al. 2016) which, in principle, enables them to resist the forces imposed by SEM preparation. When coccoliths are severely malformed, however, the interlocking is impaired and coccospheres are more likely to collapse.

**Figure 1:**
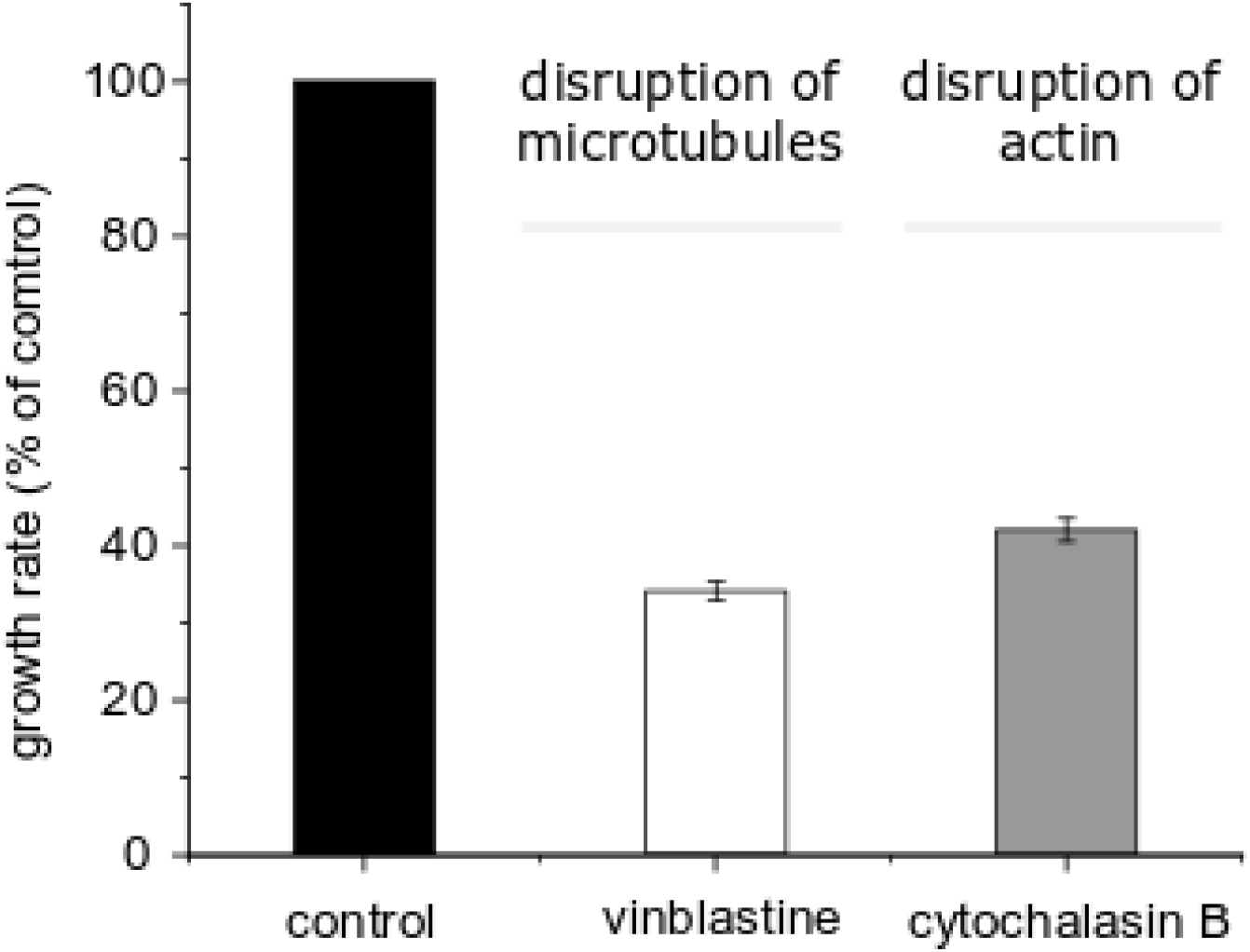
Effects of cytoskeletal inhibitors on growth of *C. braarudii*. A) Growth rate of *C. braarudii* following treatment with 1.25 μM vinblastine or 2 μM cytochalasin B. Growth is shown relative to control (untreated) cultures as the vinblastine and cytochalasin B treatments had separate controls (specific growth rates of the controls ranged from 0.5-0.6 d^-1^). n = 3 cultures for each treatment. Error bars represent SD.

**Figure 2:**
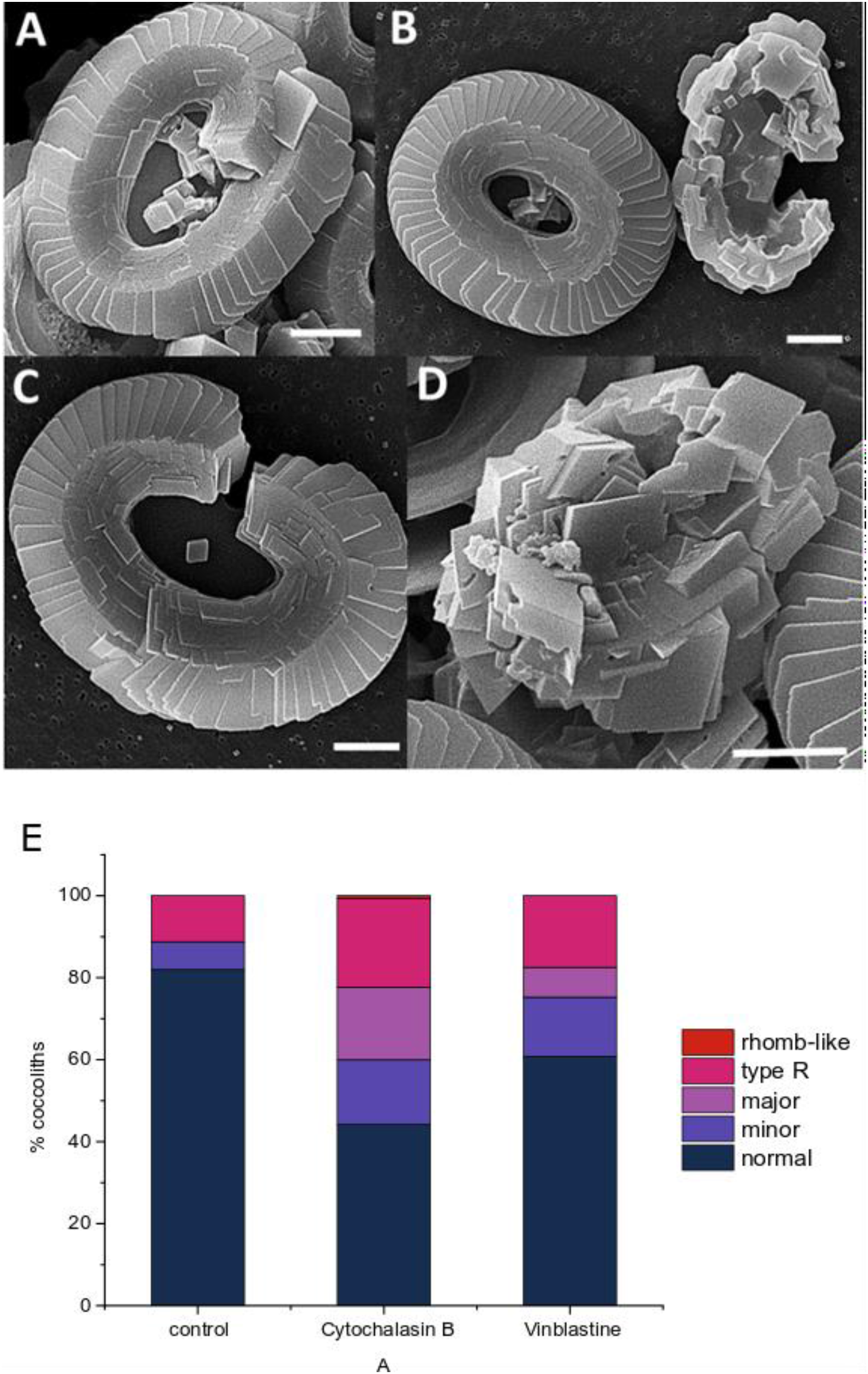
Scanning electron micrographs of the morphological categories of *C. braarudii* coccoliths. Representative SEM images of the categories used to quantify coccolith morphology. A) minor malformation, shields largely intact but elements imperfect, B) normal coccolith (left) and major malformation (right) where shields are not correctly formed. C) type R, coccolith largely intact but the shields do not form a complete ring D) rhomb-like malformation, shields are not discernible, composed of ‘blocky’ calcite crystals. Scale bars 2 μm. E) Quantification of coccolith morphology. Bars from bottom up represent the morphological categories (% of counted): normal, minor malformation, major malformation, type R and rhomb-like malformation. A minimum of 300 coccoliths were assessed for each sample, with values representing means of triplicate treatments.

**Figure 3:**
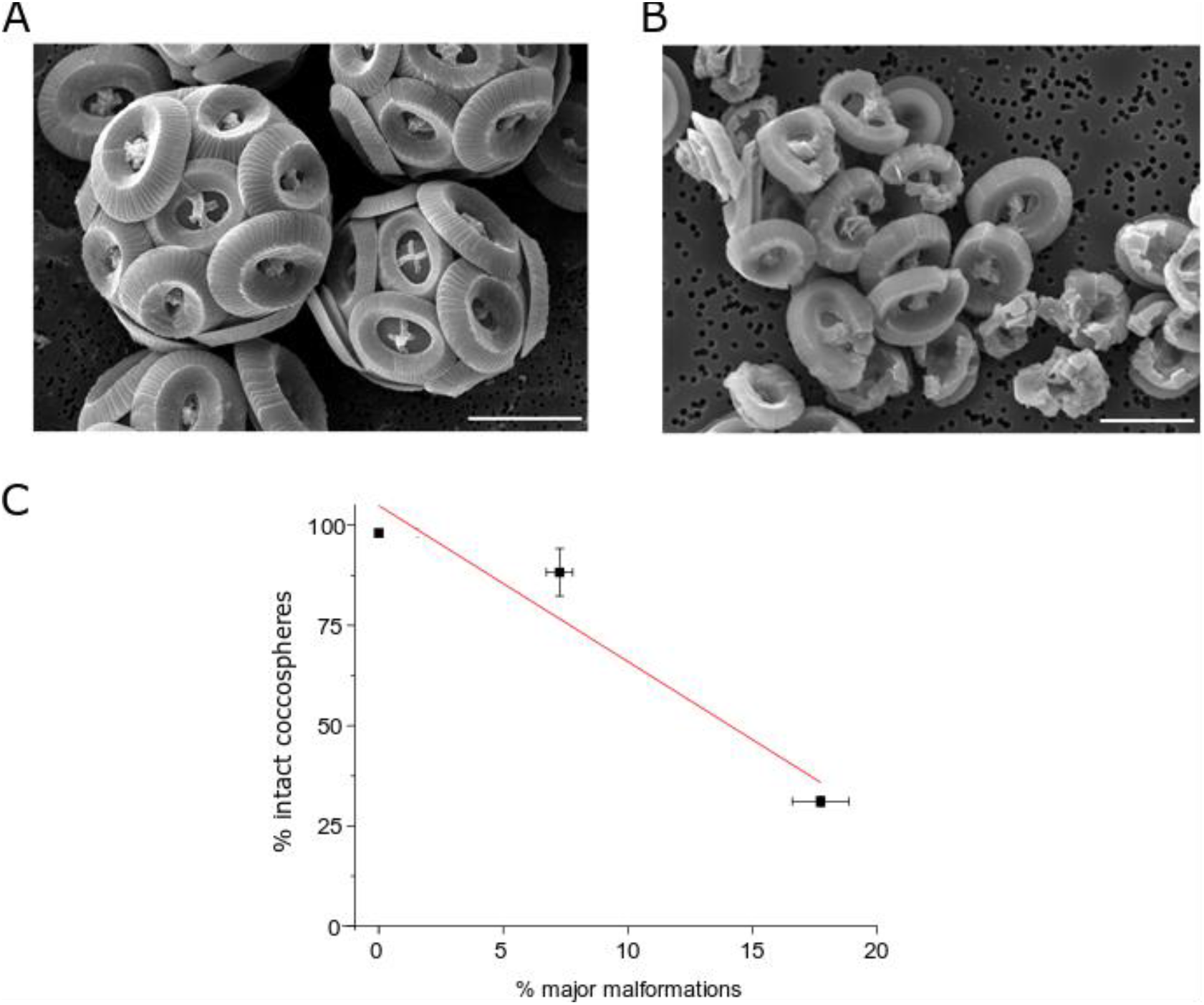
Effect of coccolith malformations on the *C. braarudii* coccosphere. **A)** SEM image of control cells showing intact coccospheres. Bar = 10 μm. **B)** SEM image from cells treated with 2 μM cytochalasin B showing collapsed coccospheres. Bar = 10 μm. **C)** Percentage of intact *C. braarudii* coccospheres versus percentage of major malformations in coccoliths. An increase in the proportion of major malformations correlates with a decrease in the % of intact coccospheres. Data points represent different treatments (control, vinblastine and cytochalasin B). The trendline represents linear regression with an r^2^ of 0.92. n= 3 replicate treatments. Error bars represent SD.

**Figure 4:**
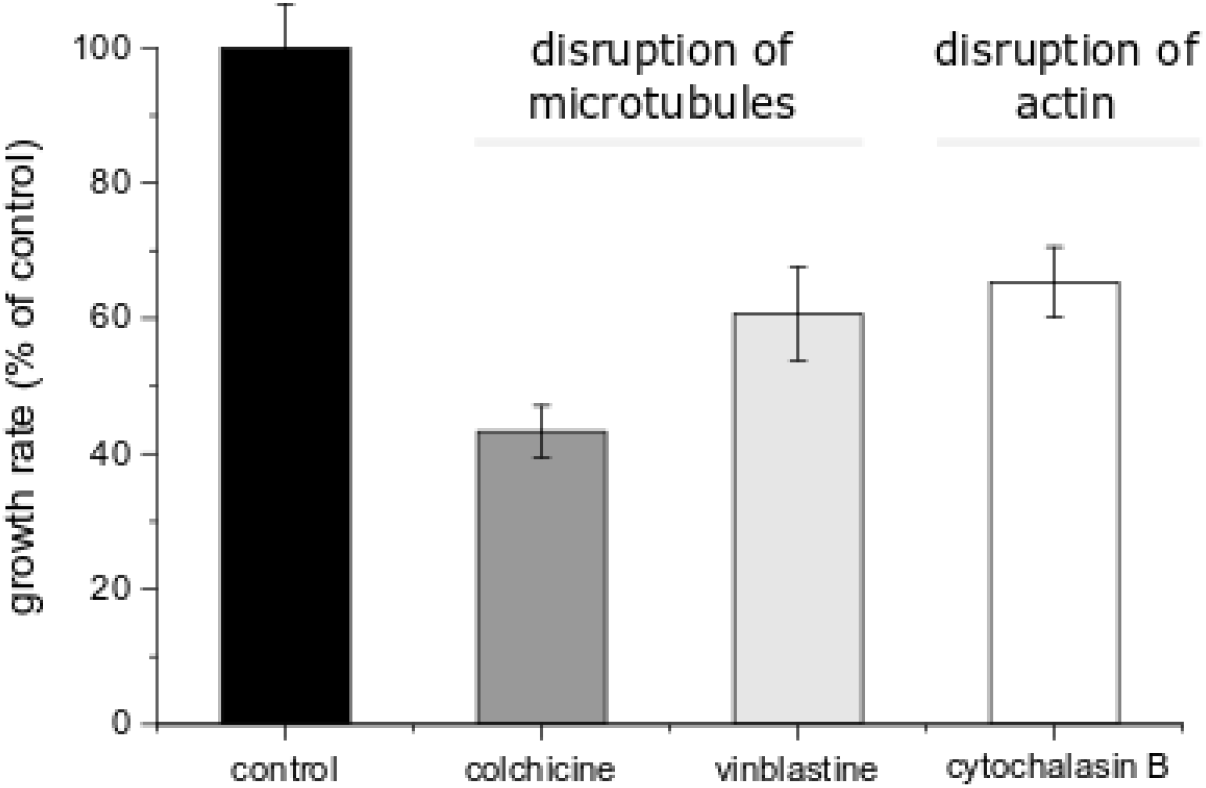
Effects of cytoskeletal inhibitors on growth of *S. apsteinii*. Specific growth rate of *S. apsteinii* (shown relative to the control) following treatment with 20 μM colchicine, 1.25 μM vinblastine or 1 μM cytochalasin B. n = 3 cultures for each treatment. Error bars represent SD.

**Figure 5:**
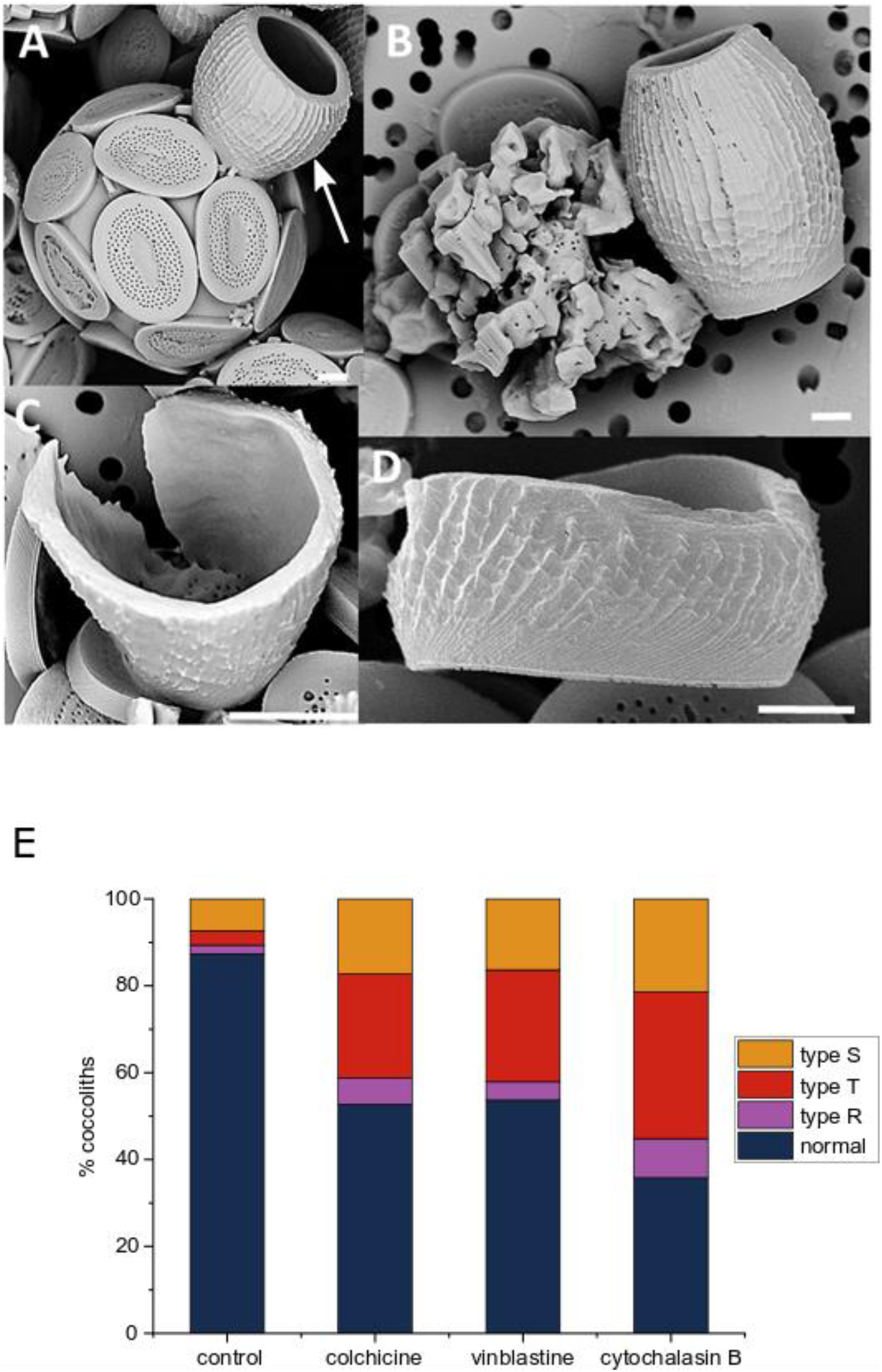
Scanning electron micrographs of the morphological categories of *S. apsteinii* coccoliths. Representative SEM images of the categories used to quantify coccolith morphology. A) Intact coccosphere with disc-like muroliths and barrel-shaped lopadolith (arrowed) exhibiting normal morphology, B) type T (heavily malformed, loss of barrel shape) (left) and normal (right), C) type R, lopadoliths barrel formed normally except that it does not form a closed cylinder. D) type S, short lopadolith with no obvious teratological malformation. Coccoliths shown in B, C, and D are lopadoliths. Scale bars 2 μm in A, B, D, and 5 μm in C. E) Quantitation of coccolith morphology in *S. apsteinii*. Bars from bottom up represent the morphological categories (% of counted): normal, type R, type T, type S. A minimum of 300 coccoliths were assessed for each sample, with values representing means of triplicate treatments.

## Results

### Effects of cytoskeletal inhibitors on *C. braarudii*

To examine the impacts of disrupting the cytoskeleton on coccolith formation in *C. braarudii*, we treated cells with the microtubule inhibitor vinblastine and the actin inhibitor cytochalasin B. As the cytoskeleton is essential for cell division and many other cellular processes, a total disruption of cytoskeletal function would prevent cell growth or secretion of coccoliths. We therefore performed a series of pre-experiments to determine inhibitor concentrations that allow the cells to continue to grow at a reduced growth rate, indicating that the inhibitor disrupts the cytoskeleton to some extent but does not completely impair secretion or cell division.

Growth of *C. braarudii* cells in 2 μM cytochalasin B to disrupt actin networks resulted in a 58% reduction in growth rate (Fig 1). Application of 1.25 μM vinblastine to disrupt microtubule function reduced growth by 66%. Scanning electron microscopy (SEM) was then used to examine coccolith morphology in these cultures. Despite the similar reduction in growth rate, the effects of the two different inhibitors on coccolith morphology were markedly different. In general, the effects of cytochalasin B were more severe than the ones of vinblastine. In particular, the percentage of major malformations (0.0% in the control) rose to 7.3 ± 0.5 % under the influence of vinblastine but to 17.7 ± 1.1 % under cytochalasin B treatment (n=3, ± SD) (Fig 2). The level of minor malformations did not differ between the two inhibitors.

The effects on coccolith morphology following cytoskeletal disruption was reflected in the percentage of intact coccospheres present during SEM analysis. Whilst control cells displayed 98.1 ± 0.9 % intact coccospheres, only 88.3 ± 5.9 % of coccospheres were intact after treatment with 1.25 μM vinblastine and only 31.1 ± 1.2 % after treatment with 2 μM cytochalasin B (n=3, ± SD) (Fig 3A-C). Since minor malformations by definition do not affect the double shield architecture that is instrumental in forming an interlocking coccosphere, coccoliths displaying minor malformations are still able to integrate normally into the coccosphere. Coccoliths displaying major malformations, by contrast, do not interlock and therefore make the coccosphere unstable. This is reflected in the correlation between intact coccospheres and major malformations (Fig 3C). The dependence of intact coccospheres on coccolith morphology was also observed in *Calcidiscus leptoporus* but was not quantified (Langer et al. 2006, Langer and Bode 2011). The relationship between coccolith morphology and coccosphere integrity highlights the importance of coccolith morphogenesis in coccolithophore ecology and evolution. The significance of an intact coccosphere has at least two aspects. First, an interlocking coccosphere has remarkable mechanical properties (Jaya et al. 2016) which will be impaired by heavily malformed coccoliths. Second, *C. braarudii* cannot grow without a coccosphere (Walker et al. 2018).

### Effects of cytoskeletal inhibitors on *S. apsteinii*

We grew *S. apsteinii* in the presence of the microtubule inhibitors vinblastine and colchicine, and the actin inhibitor cytochalasin B. Treatment with 1.25 μM vinblastine and 20 μM colchicine for reduced the growth rate by 39% and 57% respectively (Fig 4). Treatment with 1 μM cytochalasin B for ca 20 days reduced the growth rate by 35%. The inhibitor concentrations used to cause a moderate reduction in growth rate in *S. apsteinii* are therefore similar to those in *C. braarudii*. However, these concentrations are markedly different from *E. huxleyi* (Langer et al. 2010) (Supplementary Table 1), which may point to differences between species in the types of actin and microtubules (Thompson et al. 1984, Gunning et al. 2015, Howes et al. 2018) or in the uptake of the inhibitors.

The cytoskeleton inhibitors also had pronounced effects on coccolith morphology in *S. apsteinii*. We did not quantify the effects of cytoskeletal disruption on murolith morphology, although a qualitative analysis indicated that murolith malformations increased under all tested inhibitors (Supplementary Figure 1). Quantitative analysis of lopadolith morphology revealed a similar morphological response to all tested inhibitors (Fig 5). The proportion of Type S, type T and type R malformations increased in under all treatments, with the proportion of normal coccoliths decreasing from 87% in control cultures to 36-54% in the presence of cytoskeleton inhibitors. As the nature of the malformations are similar between all treatments, we conclude that disruption of either the microtubule or actin networks have similar effects on coccolith morphology. It is interesting nonetheless that the treatment that had the greatest impact on growth (colchicine) did not have the greatest impact on morphology (cytochalasin B).

The coccoliths of *S. apsteinii* do not interlock, unlike those found on *C. braarudii* and *E. huxleyi*, and so do not usually remain intact during sample preparation for SEM analysis. We therefore did not quantify collapsed coccospheres.

## Discussion

Our results suggest that both microtubules and actin filaments are involved in coccolith morphogenesis in *C. braarudii* and *S. apsteinii*, in support of previous findings in *E. huxleyi* (Langer et al 2010). Unlike the application of the silicon analogue germanium, which results in distinct types of malformed coccoliths in *C. braarudii* and *S. apsteinii* (Langer et al 2021), the malformations induced by disruption of the cytoskeleton were not specific to this stress (Giraudeau et al. 1993, Langer et al. 2006, Benner 2008, Gerecht et al. 2015). Although cytoskeletal inhibitors did not cause specific malformations, the effects on coccolith morphogenesis are unlikely to be simply a result of general cellular stress. Disruption of coccolith formation did not simply correlate with inhibition of growth, as treatments that gave the greatest inhibition of growth (e.g. colchicine to *C. braarudii)* did not result in the highest degree of coccolith malformations (Fig 3 & 5). This also supports observations from *E. huxleyi* that growth inhibition via other mechanisms, such as the inhibition of photosynthesis, do not result in an increase in coccolith malformations (Langer 2010). We therefore propose that the malformations we observe point to a requirement for both actin and microtubules in shaping the developing coccolith.

Disruption of actin networks with cytochalasin B in *C. braarudii* or *S. apsteinii* did not have a distinct effect on coccolith morphology from the disruption of microtubules with either colchicine or vinblastine, suggesting that both components of the cytoskeleton contribute to similar aspects of coccolithogenesis. This finding differs from an earlier study where disruption of actin with latrunculin B in *C. braarudii* inhibited coccolith secretion, suggesting a specific role for actin in this process (Durak 2017). There are several explanations for these differing results. The phenotypic difference may simply reflect a difference in the effective concentration of the inhibitor applied, as it is difficult to gauge the extent to which the actin network has been disrupted in the two studies. The differing phenotypes may also result from the differences in the mode of action of latrunculin B and cytochalasin B. While the former depolymerizes actin the latter caps actin filaments thereby reducing polymerization rate (MacLean-Fletcher and Pollard 1980, Forscher and Smith 1988, Gibbon et al. 1999, Foissner and Wasteneys 2007). Differences in the experimental protocols may also have contributed to the different phenotypes. Whilst the current study observed coccolith production over several generations, Durak et al 2017 observed production of new coccoliths 24 h after decalcification. Whilst this has the advantage of ensuring that only coccoliths produced following the application of the treatment are observed, the process of decalcification itself may induce malformations (unpublished observations G. Langer 2017)

Durak et al. (2017) observed only a few heavily malformed coccoliths in SEM samples from *C. braarudii* cultures treated with latrunculin B and hypothesised that these arose from intracellular coccoliths that had not been secreted. The nature of these malformations differ from those observed in response to cytochalasin B or in response to other stressors (Giraudeau et al. 1993, Langer et al. 2006, Benner 2008, Gerecht et al. 2015). It is therefore possible that the decalcification process contributed to these unusual malformations. In support of this conclusion, similar malformations were also observed in recalcifying cells in response to the microtubule inhibitor nocodazole (17 μM) (Durak 2017), but were not seen in the current study following treatment with 2 μM vinblastine. Again, these phenotypic differences could be due to differences in the concentration of inhibitor applied or their mode of action. Both nocodazole and vinblastine stabilize microtubule ends at nanomolar concentrations but depolymerize them at micromolar concentrations (Jordan et al. 1992). A difference in their effect on coccolith morphogenesis could therefore stem from the different concentrations used. However, given the relatively high proportion of malformations in recalcifying control cells, it is likely that the unusual malformations observed in response to nocodazole are the result of a combined effect of decalcification and nocodazole (Durak et al. 2017).

Disruption of the cytoskeleton could potentially influence the calcification processes in multiple ways, from the intracellular transport of substrates to the coccolith vesicle, to the direct shaping of the coccolith vesicle and the exocytosis of the mature coccolith (Durak 2017). Disruption of the morphogenetic role of the cytoskeleton implies that cytoskeleton inhibitors should cause teratological malformations, rather than incomplete growth (Young and Westbroek 1991). Incomplete coccoliths may arise if transport of substrates to the coccolith vesicle is disrupted, or if the cytoskeleton is involved in the cellular ‘stop signal’ that prevents further crystal growth following the formation of a fully mature coccolith. In the present study, we found little evidence for the presence of incomplete coccoliths following disruption of the cytoskeleton in *C. braarudii*. We did not quantify incomplete coccoliths in *C. braarudii* because a preliminary analysis revealed that the percentage of incomplete coccoliths in all samples was less than 1%. The presence of incomplete coccoliths in *S. apsteinii* is slightly more difficult to resolve because it is not entirely clear whether the S-type malformation should be classified as incomplete, malformed (in the strict teratological sense), or normal. The S-type category is a short lopadolith (i.e. the length of the barrel is reduced) with no obvious teratological malformation. While this might seem to suggest that it should count as incomplete, there are several observations from different cultures suggesting that there is a great variability in lopadolith size including S-type-size (not quantified). Given this variability, it is possible that we should consider the S-type morphology as normal, rather than a malformation. However, as the S-type morphology is more abundant in response to cytoskeletal inhibitors, it does appear to represent a genuine effect, albeit an effect on size rather than “completeness”. The distinction between size and incompleteness is harder to define in *S. apsteinii* than in *E. huxleyi*, in which incomplete coccoliths can be clearly identified by the absence of a well-defined rim while complete coccoliths can exhibit different sizes (Langer et al. 2010, Rosas-Navarro et al. 2016). The example of *E. huxleyi* shows that, from a mechanistic point of view, there is a distinction between size and incompleteness. An incomplete coccolith represents a situation where crystal growth was stopped too early, so that the coccolith does not possess all the normal structural features, i.e. the cellular “stop-signal” was not given correctly. This situation is therefore distinct from a normal coccolith of small size. In terms of *S. apsteinii*, this means that the increase in the S-type morphology in response to cytoskeleton inhibitors does not indicate a connection between the “stop-signal” for coccolith growth and the cytoskeleton. The absence of an increase in incomplete coccoliths in *C. braarudii* (this study) or *E. huxleyi* (Langer et al. 2010) further suggests that cytoskeleton inhibitors applied in this manner do not affect the stop-signal for coccolith growth.

In summary, our findings show that both actin filaments and microtubules are involved in coccolith morphogenesis in *C. braarudii* (Coccolithales) and *S. apsteinii* (Zygodiscales). Taken together with previous findings in *E. huxleyi* (Isochrysidales) data (Langer et al. 2010), this strongly suggests that these two cytoskeleton elements play a central role in coccolith morphogenesis in coccolithophores. Detailed examination of the mechanisms through which the actin and microtubules interact to influence the shape of the coccolith vesicle as the coccolith matures is now required to fully test the ‘dynamic mould’ hypothesis. To achieve this novel high-resolution imaging techniques, which preserve sub-cellular features, such as cryo-FIB-SEM will likely be helpful. These highly specialised electron microscopy applications are difficult and time consuming, but the results of this study show that the effort is worthwhile. Our data suggest that the cytoskeleton is at the heart of coccolith morphogenesis.

## Acknowledgements

The work was supported by an NERC award to GLW (NE/N011708/1), an NSF-GEO award to ART (1638838), and an ERC Advanced Grant to CB (ERC-ADG-670390). Electron microscopy analyses were performed at the PEMC (Plymouth University, UK). The authors declare no competing interests.

## Supplementary Information

**Supplementary Figure 1:**
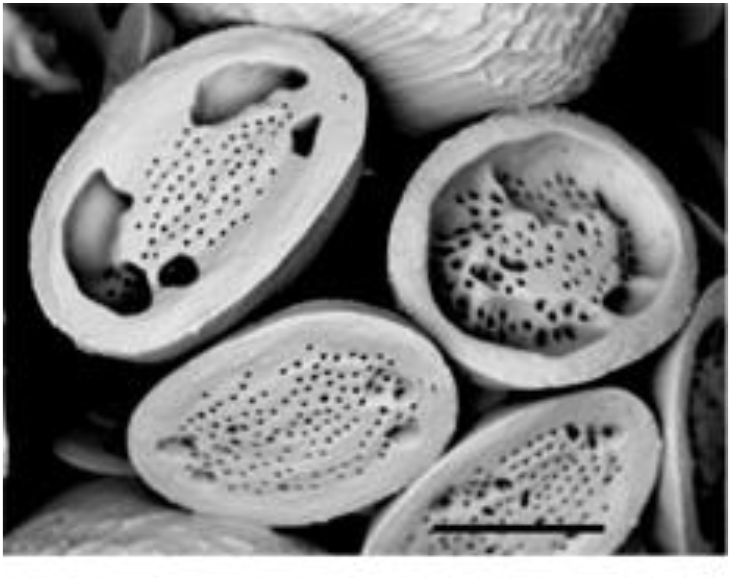
Cytoskeleton inhibitors also induce malformations in *S. apsteinii* muroliths. In addition to barrel-shaped lopadoliths, *S. apsteinii* also produces disc-like muroliths (Figure 5). Muroliths also exhibited distinct malformations in cells treated with cytoskeleton inhibitors, although these were not quantified. The SEM image shown illustrates malformed muroliths from a *S. apsteinii* cell treated with 1 μM cytochalasin B. Bar = 4 μm.

**Supplementary Table 1:** Summary of cytoskeleton inhibitors used to disrupt coccolithophore calcification.

| Species | Cytoskeleton inhibitor | Mode of action | Concentration used |
| --- | --- | --- | --- |
| **Microtubule inhibitors** |  |  |  |
| *E. huxleyi*<br>(Langer 2010) | Colchicine | Polymerisation inhibitor<br>(colchicine domain) | 1000 µM |
| *C. braarudii*<br>(Durak 2017) | Nocodazole | Polymerisation inhibitor<br>(colchicine domain) | 17 µM |
| *C. braarudii*<br>This study | Vinblastine | Polymerisation inhibitor<br>(vinca domain) | 2 µM |
| *S. apsteinii*<br>This study | Colchicine | Polymerisation inhibitor<br>(colchicine domain) | 20 µM |
| *S. apsteinii*<br>This study | Vinblastine | Polymerisation inhibitor<br>(vinca domain) | 1.25 µM |
| **Actin inhibitors** |  |  |  |
| *E. huxleyi*<br>(Langer 2010) | Cytochalasin B | Prevents polymerization | 100 µM |
| *C. braarudii*<br>(Durak 2017) | Latrunculin B | Prevent polymerization<br>enhance depolymerisation | 1 µM |
| *C. braarudii*<br>This study | Cytochalasin B | Prevents polymerization | 1.5 µM |
| *S. apsteinii*<br>This study | Cytochalasin B | Prevents polymerization | 1 µM |

